# High throughput single-cell proteomics of in-vivo cells

**DOI:** 10.1101/2024.11.01.621461

**Authors:** Shiri Karagach, Joachim Smollich, Ofir Atrakchi, Vishnu Mohan, Tamar Geiger

## Abstract

Single-cell mass spectrometry-based proteomics (SCP) can resolve cellular heterogeneity in complex biological systems and provide a system-level view of the proteome of each cell. Major advancements in SCP methodologies have been introduced in recent years, providing highly sensitive sample preparation methods and mass spectrometric technologies. However, most studies present limited throughput and mainly focus on the analysis of cultured cells. To enhance the depth, accuracy, and throughput of SCP for tumor analysis, we developed an automated, high-throughput pipeline that enables the analysis of 1,536 single cells in a single experiment. This approach integrates low-volume sample preparation, automated sample purification, and LC-MS analysis with the Slice-PASEF method. Integration of these methodologies into a streamlined pipeline led to a robust and reproducible identification of more than 3000 proteins per cell. We applied this pipeline to analyze tumor macrophages in a murine lung metastasis model. We identified over 1,800 proteins per cell, including key macrophage markers and ∼500 differentially expressed proteins between tumor and control macrophages. PCA analysis successfully separated these populations, revealing the utility of SCP in capturing biologically relevant signals in the tumor microenvironment. Our results demonstrate a robust and scalable pipeline poised to advance single-cell proteomics in cancer research.

## Introduction

Analyzing biological systems at the single-cell level offers the advantage of capturing pure cellular signals, eliminating the need to average signals across predefined cell populations based on known markers. The advent of single-cell RNA sequencing (scRNA-seq) has revolutionized our ability to identify novel cell populations and trace cellular trajectories, significantly advancing our understanding of developmental processes and dynamic cellular changes across various biological systems. In cancer research, scRNA-seq has been instrumental in identifying unique cell populations that may influence therapeutic responses— insights often masked in population-level analyses due to the averaging of signals, which obscures the presence of rare cell types(1, 2).

Despite its transformative impact, scRNA-seq has limitations. RNA-level analyses alone cannot fully capture cellular function, as RNA and protein expression are not always correlated(3, 4). Numerous studies, including ours, highlight the importance of protein-level analyses for understanding cellular pathways(3, 5). Pathways involving complex molecular machinery, such as the ribosome, spliceosome, electron transport chain, and proteasomes, often display low RNA-protein correlation. In contrast, pathways under transcriptional regulation, such as the interferon pathway, exhibit high RNA-protein correlations. Signaling, metabolic, and adhesion pathways typically show moderate correlations (around 0.5), though these correlations vary widely across biological systems (6-9). Additionally, scRNA-seq is often limited by low transcriptional coverage per cell, high variability in signal between cells, and significant amounts of missing data(10). As a result, rigorous statistical analyses generally require data from thousands of cells to define well-resolved cellular clusters with sufficient depth.

The recognition of scRNA-seq’s capabilities and limitations has spurred the development of single-cell proteomics (SCP) technologies to broaden the scope of mass spectrometric analyses. Advances in mass spectrometry sensitivity, driven by improved ion transfer efficiency and reduced background noise, now permit the analysis of sub-nanogram protein quantities (11-17). Enhanced sample preparation techniques, such as minimizing protein absorption to plasticware and reducing sample volumes, improve protein recovery and digestion efficiency by increasing protein concentration (18-21). To date, most SCP studies have focused on single cultured cells, demonstrating advancements in each of these parameters. In this study, we aim to further advance SCP technology for the analysis of tumor cells. We developed an automated sample preparation protocol to enable high-throughput single-cell processing in 1536-well plates. By incorporating cell fixation, we established a robust and scalable pipeline essential for single-cell tumor proteomics.

## Experimental Procedures

### Cell culture

HeLa cells and MC38-RFP cells were cultured in DMEM supplemented with 10% FBS and 1% Penicillin-Streptomycin at 37°C with 5% CO2. To generate single-cell suspensions, cells were dissociated from the culture plates with trypsin, washed with PBS, and diluted to a concentration of 100-200 cells/μl in degassed PBS for downstream single-cell separation.

### Mouse models and macrophage isolation

MC38 cells were injected into the tail vein of 10-week-old C57BL6 mice (Harlan). Mice were injected with 10^6 cells in 100 μl PBS or with the same volume of PBS as control. One week after injection, mice were euthanized, and lungs were removed and dissociated into single cells using the lung dissociation kit (Miltenyi) on the gentleMACS Dissociator (Miltenyi). Red blood cells were removed using the red blood cell lysis buffer. Cells were stained with Live/dead Fixable Blue (L34961, Thermo Fisher Scientific) to remove dead cells and then labeled with the following antibodies to enable downstream macrophage sorting: CD45 (#103107 Biolegend), CD11b (#740861 BD-Biosciences), CD11c (#565452, BD-Biosciences), CD64 (#139332 Biolegend). Isolation of pooled macrophages was performed by flow cytometry on the FACS-Aria III sorter (BD-Biosciences).

### Single-cell dispensing on CellenOne and in-solution trypsin digestion

Single cells were isolated using the CellenOne automated image-based single-cell dispenser. This instrument utilizes piezoelectric technology for gentle, picoliter-volume acoustic dispensing, which enhances cell viability and allows for the miniaturization of sample preparation workflows. Uncoated Piezo Dispensing Capillaries (PDC) of size M or L were used in all experiments. The lysis and digestion buffer MasterMix included 0.2% n-dodecyl-β-D-maltoside (DDM), 20 ng/μL Sequencing Grade Modified Trypsin (Promega), and 100 mM Triethylammonium bicarbonate (TEAB) buffer. A polystyrene 384-well plate served as the probe, and polypropylene 384-well plates (Eppendorf) or 1536-well plates (Greiner) were used as the target. To ensure droplet stability, the intermediate task DipDropCheck was performed every 14 wells during MasterMix dispensing. Relative humidity was maintained at 50%, and the temperature was regulated using a DewPoint set point to prevent condensation. After dispensing, the plates were covered and centrifuged for 1 minute at 800g. The single-cell suspensions were prepared from tissue-cultured HeLa cells or dissociated mouse lungs, diluted into a final solution of 100-200 cells/μL suspension in degassed PBS. To ensure homogeneous sampling during the isolation process, the Mix&Take task was used to prevent cell settling. Based on cell mapping prior to isolation, diameters of 20-35 μm and 15-30 μm were used for HeLa cells and macrophages, respectively. A maximum elongation of 1.85 was applied to avoid the isolation of doublets or non-single cells. During isolation, humidity and temperature conditions matched those described for MasterMix dispensing. The plates were covered and centrifuged again for one minute at 800g. Two different incubation approaches were tested: 1) Covered plate with an aluminum seal and 2) Water rehydration at 500Hz. In both approaches, the plates were incubated at 50°C for two hours with 85% humidity to prevent evaporation. For the 1536-well plate, an in-house produced aluminum adaptor with an anti-corrosion coating, was used to improve heat conductivity. After incubation, the plates were cooled to 20°C and centrifuged again for one minute at 800g. Automated dilution was performed using 5 μL of solvent A (99.9% water + 0.1% formic acid) with the Bravo liquid handler (Agilent). Finally, the plates were frozen at -80°C or used for semi-automated Evotip loading.

#### Semi-automated Evotip loading

Sample clean-up and loading were performed based on the semi-automated Evotip loading protocol of Evosep for the Bravo liquid handler (Agilent). The publicly available protocol was adapted to the 96-ST head. For the semi-automated sample loading from 1536-well plates, the V10 10 μl tips (Agilent) were used. To avoid solvent leakage during solvent pipetting on the robot, “air gaps” (post aspiration volumes of 5ul) were added to each pipetting step. In addition, to ensure complete pipetting without any sample loss, “blow-out” options were chosen when sample solutions were dispensed.

### LC-MS analysis

LC-MS analyses were performed on the Evosep1 liquid chromatography system (Evosep) coupled to the TimsTOF SCP mass spectrometer (Bruker). Peptides were separated on the Aurora Elite CSI 15×75 C18 UHPLC column (Ionopticks) using the Whisper Zoom 40 SPD method. Mass spectrometric acquisition was performed in a data-independent (dia) PASEF mode. For the diaPASEF mode the ion accumulation and ramp time was set to 100ms and precursors in an ion mobility range from 1/K0 = 1.45 Vs cm-2 to 0.64 Vs cm-2 were analyzed over a mass to charge (m/z) range from 100 to 1700. Precursor ions for MS/MS analysis were isolated based on diaPASEF window placement covering an ion mobility range from 1/K0 = 0.75 Vs cm-2 to 1.2 Vs cm-2 and a mass to charge (m/z) range of 400 – 1000. The estimated cycle time was 0.64 s with 1 MS ramp and 5 MS/MS ramps, resulting in 75 MS/MS windows. The collision energy was ramped from 20eV at 1/K0 = 0.6 Vs cm-2 to 59eV at 1/K0 = 1.6 Vs cm-2. Singly charged precursors were excluded with a polygon filter. We used the publicly available Slice-PASEF window scheme for design (22). The method file “diaParameters” was downloaded and included in the TimsTOF SCP acquisition method file.

### Data analysis

DIA-NN version 1.9.1 was used to analyze the mass spectrometric raw files. Analyses of HeLa files were performed using a library-based approach (spectral library provided by BRUKER containing 67686 proteins and 13827 genes). Library precursors were reannotated using a FASTA database (FASTA files UP000000589_10090.fasta and UP000000589_10090_additional.fasta, version August 2022). Trypsin was selected as protease, with a maximum of one missed cleavage. DIA-NN default settings were used with the following exceptions: Precursors’ charge range was set to 2-10, and the “unrelated runs” option was selected. Precursor m/z range was set to 300 – 1700, and the “Fragment ion m/z range” was set to 200 - 1700. For Slice-PASEF samples, the command “—tims-scan” was added in the “Additional options” in DIA-NN. Statistical analysis was performed in R. For the analysis of the murine macrophage samples, we used an in-house generated spectral library based on previously acquired .dia samples (containing 29438 proteins and 10517 genes).

### Experimental Design and Statistical Rationale

All experiments analyzed single cells, which form biological replicates. All technical analyses of HeLa cells included at least 12 cells per group. These experiments were repeated at least three times, reaching similar conclusions. One representative experiment is presented in each graph. Each figure shows a separate experiment with no overlapping samples. In-vivo tumor mouse analyses were performed on three mice, with more than 70 cells analyzed per group.

### Data Availability Statement

All protein data is provided in supplementary tables included in the submission. All raw files will be uploaded to Pride.

## Results

The application of single-cell proteomics to tumor samples demands robustness, depth, accuracy, and throughput. Building on previous advancements in low-volume sample preparation, we developed an experimental pipeline that integrates automated, high-throughput sample preparation for LC-MS analysis to achieve deep proteomic coverage (**Fig. 1A**). Our objective was to enable the analysis of over 1,000 cells within a single experiment, a capability essential for *in-vivo* studies and the foundation for advancing single-cell biological research.

**Figure 1:**
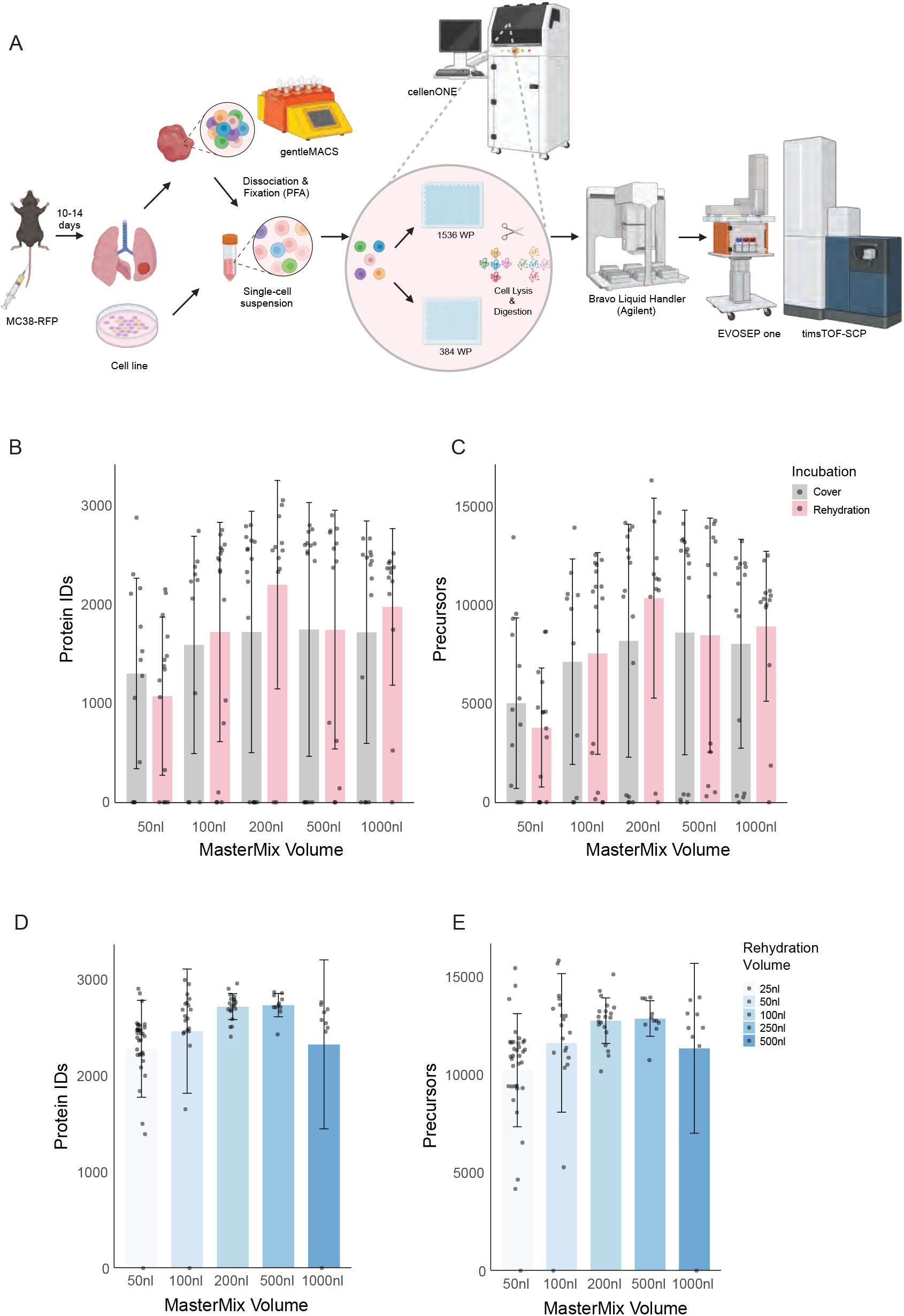
Miniaturization of SCP workflow. **A**. A schematic representation of the SCP workflow. Sample preparation is initiated with cultured cells or in-vivo tumor samples. Single-cell suspensions are separated into single wells on the CellenOne dispenser directly into MasterMix. Reactions are loaded onto Evotips on the Bravo liquid handler before LC-MS analysis on the Evosep1 couple to the TimsTOF SCP mass spectrometer. Figure is created with BioRender.com **B**. Barplot shows the number of identified protein groups using increasing MasterMix volumes, comparing covered plates vs. rehydration with 300 nL water. **C**. Plot of the number of precursors for the same experiment as B. **D**. Same as B, but with rehydration with volumes equal to 50% of the initial reaction volume. **E**. Plot of the number of precursors of the same experiment as D.

### High-throughput single-cell proteomics sample preparation

As the initial step in sample preparation optimization, we developed a protocol for single-cell separation on the CellenONE dispenser and in-solution trypsin digestion. The CellenONE system allows low-volume dispensing and reduces sample evaporation through controlled temperature and humidity. Additionally, it supports protocol execution in open plates while continuously rehydrating the wells. Following standard protocols, we assessed proteomic coverage by preparing samples in increasing volumes of MasterMix within a 384-well plate. In the first set of experiments, plates were covered to minimize evaporation. In the second set, we rehydrated each well with 300 nL of water during a 2-hour digestion. Samples were subsequently loaded onto Evotips and analyzed on the Tims TOF SCP. Our analyses found a consistent improvement in protein identification when using the rehydration protocol compared to covered wells (**Figure 1B, C, Supplementary Table S1**). Furthermore, we observed that reducing the sample volume below 100 nL significantly reduced proteomic depth, likely due to increased evaporation. Notably, proteome coverage showed only minor differences upon dehydration, with mean coverage ranging from 1800-2000 proteins and maximal coverage between 2900-3100 proteins, independent of the initial MasterMix volume. We speculated that the large 300 nL rehydration volume may have masked initial volume differences. We then repeated the experiments with rehydration volumes proportional to the initial reaction volumes. Specifically, rehydration with 50% of the initial reaction volume improved coverage, particularly for smaller reaction volumes of 50, 100, and 200 nL. Reactions using 50 nL were suboptimal, but reaction volumes of 100-200 nL, combined with rehydration of 50-100 nL, provided optimal results in a 384-well plate format (**Figure 1D, E**).

We automated the Evotip loading protocol on a Bravo liquid handler (Agilent) to further increase throughput and reproducibility. This protocol allows for the automated loading of 96 Evotips from 384-well- or 1536-well plates while minimizing sample loss (**Figure 2A**). A comparison of 40 single HeLa cells showed low variation in the number of peptides and proteins identified per cell (mean protein count 3101 ± 203 standard deviation, **Figure 2B, C, Supplementary Table S2**). Determination of the coefficient of variation (CV) for all detected proteins showed a median CV of 24.9%, indicating high quantitative accuracy and robustness (**Figure 2D**).

**Figure 2:**
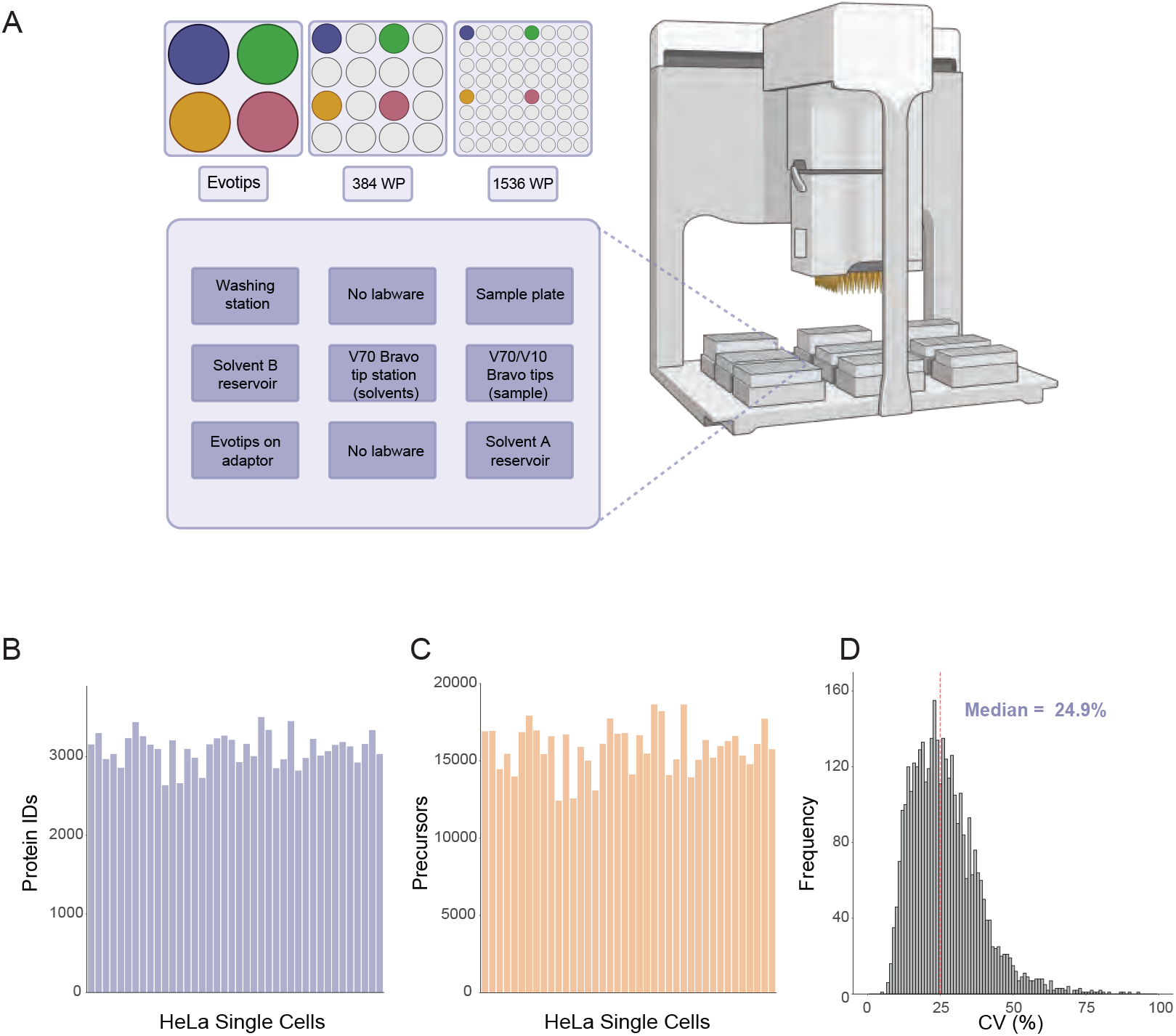
Semi-automated Evotip loading. **A**. A semi-automated Evotip loading protocol on the Bravo liquid handler enables sample loading from 384-well and 1536-well plates onto 96-well format Evotips. **B-C**. Automated loading of 40 HeLa cells results in reproducible protein (**B**) and precursor (**C**) identification. **D**. Determination of the coefficient of variation of each protein shows overall low CVs across 40 samples.

With the automated pipeline in place, we explored options to increase further sample preparation throughput—critical for single-cell proteomics in tumor analysis, where thousands of cells must be analyzed to define cellular clusters in an untargeted manner. To achieve this, we transitioned from 384-well to 1536-well plates, allowing reduced sample volumes while limiting rehydration due to the small well size. To accommodate the CellenONE system for this format, we designed a metal adaptor that fits the height of a 1536-well plate and facilitates temperature control during sample preparation (**Figure 3A**). Testing MasterMix volumes ranging from 20 to 200 nL, we observed the highest peptide and protein identification rates with 100 nL volumes (**Figure 3B, C, Supplementary Table S3**). Direct comparisons between the two plate formats showed significantly greater proteome coverage in 1536-well plates versus 384-well plates for 50-100 nL reaction volumes (**Figure 3D, E**). Thus, we propose that optimal reaction volumes are 200 nL for 384-well plates and 100 nL for 1536-well plates. We recommend the 1536-well format for complex tissue samples to process large cell numbers simultaneously, minimizing potential proteomic changes due to delays in cell dispensing and reducing batch effects.

**Figure 3:**
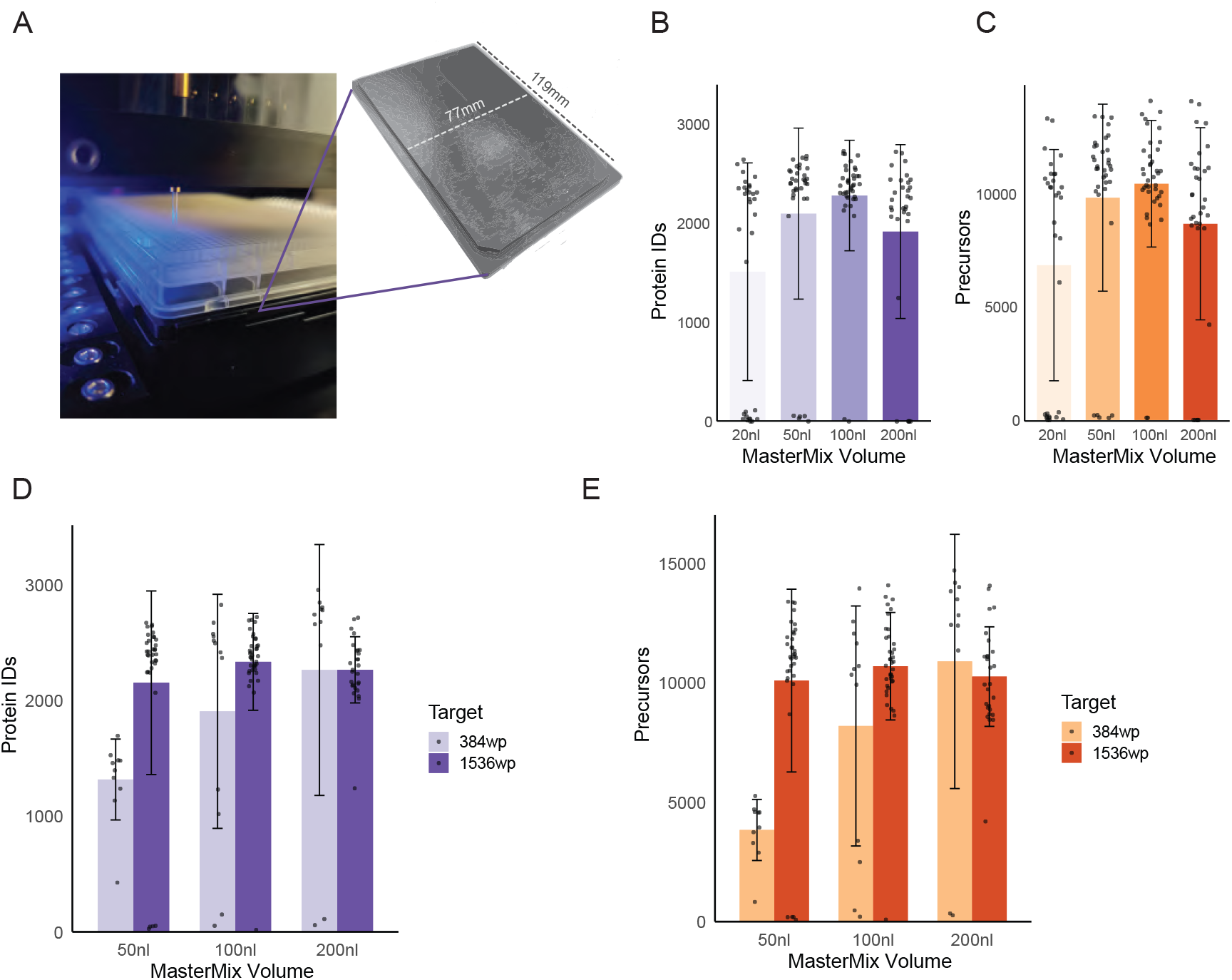
SCP on 1536-well plate format. **A**. We designed and produced an aluminum adapter for heat conductance in the CellenOne dispenser. The Piezo Dispense Capillary (PDC) is located right above the well to ensure droplet entry into the wells. **B-C**. Optimization of sample volume shows the highest number of protein groups (**B**) and precursors (**C**) when using 100 nL MasterMix volume. **D-E**. A comparison between 384-well to 1536-well format shows improved proteome (**D**) and precursor (**E**) coverage in 1536-well plates when using 50-100 nL reactions and similar results when using 200 nL volumes.

### Development of a fixation protocol for single-cell proteomics

Cell fixation can alleviate the negative impact of long gaps between cell suspension and lysis. Delayed cell lysis may lead to protein degradation and activation of stress signals that do not reflect the true biological state of the cells before cell suspension. Previous proteomic studies of archived tissue samples showed that formalin and formaldehyde have a minor impact on the proteins (23, 24), thereby allowing for robust analysis of fixed cells, however fixation impact was not yet shown for single cells. To optimize a protocol for single-cell fixation, we compared the proteomic results obtained with increasing concentrations and incubation time of paraformaldehyde (PFA) and compared them to non-fixed cells. Initial testing of cells taken immediately after fixation showed only a minor reduction in protein identification from HeLa cells fixed with 0.5% PFA for 5 and 15 minutes (**Figure 4A, B, Supplementary Table S4**). Higher concentrations and longer incubation times already decreased protein identification, presumably due to over-fixation and protein cross-linking. We then examined protein identification after overnight storage. Adding a pause point after tissue disaggregation and before single cell dispensing may be required in lengthy sample preparation procedures, and therefore it is required to ensure that cells and proteins are retained minimally altered. We incubated cells with PFA for varying durations, washed them with PBS, and kept them overnight at 4°C. Similar to the same-day measurements, we found that 0.5% PFA incubation for 15 min resulted in only a minor reduction in protein identification from single cells (**Figure 4C, D**). Surprisingly, we found no reduction in protein identification in non-fixed cells despite the overnight incubation. To deepen our investigation of proteomic changes upon fixation, we compared the proteomes of non-fixed single HeLa cells with and without an overnight incubation at 4°C. We found 153 significantly changing proteins upon incubation overnight (**Figure 4E**), suggesting that fixation is necessary in order to retain the proteomic state. In contrast, when comparing the proteomes of fixed to non-fixed cells and fixed before and after an overnight incubation, we found only 1-2 significantly changing proteins (**Figure 4F, G**). Therefore, we conclude that mild fixation retains the protein levels while only slightly affecting overall coverage. This way, fixation allows the experiments to pause without impacting biological results.

**Figure 4:**
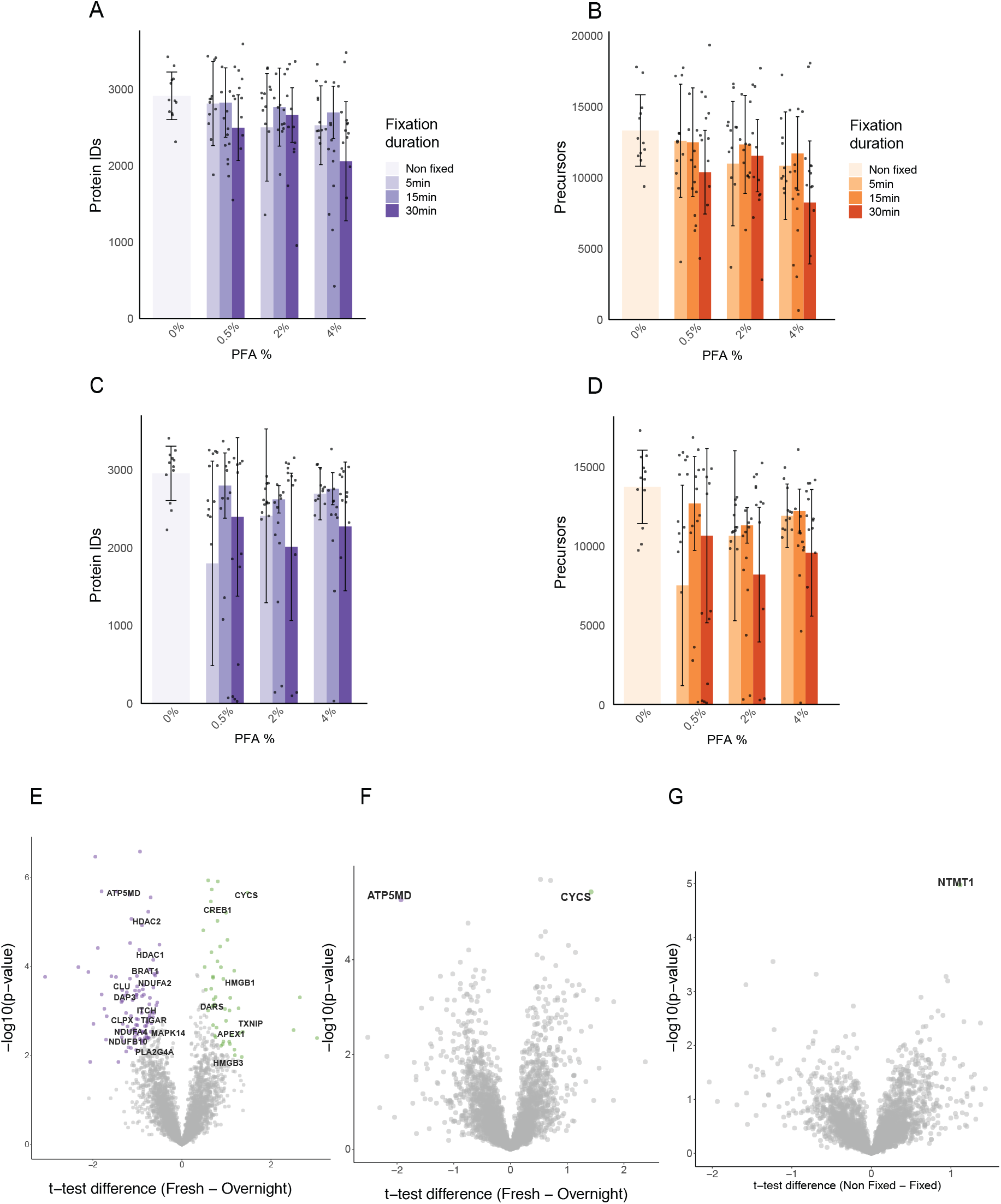
SCP of fixed cells. Cells were fixed with increasing concentration of PFA and for increasing incubation times. **A-B**. Protein (**A**) and precursor (**B**) identification when cells were dispensed and lysed on the same day, immediately after fixation. **C-D**. Protein (**C**) and precursor (**D**) identification when cells were dispensed after an overnight incubation. **E**. Volcano plot shows the significantly changing proteins in non-fixed cells, between cells before and after an overnight incubation. **F**. Volcano plot shows two significantly changing proteins in fixed cells between cells before (fresh) and after overnight incubation. **G**. Volcano plot shows no significantly changing proteins between fixed and non-fixed cells before an overnight incubation. **E-G**. T-tests were conducted with a permutation-based FDR cutoff of 5% and S0=0.1.

### Optimization of LC-MS acquisition methods

The sample preparation procedures described above provide the pipeline to prepare 1536 samples in 1-2 days. Next, we examined the ability to improve LC-MS acquisition. Our experimental setup used the Evosep1 liquid chromatography system, which provides reproducible and accurate LC parameters. We used the Whisper Zoom 40 samples per day (SPD) method. To increase the throughput, we also tested shorter methods for the acquisition of 80 and 120 samples per day (80 SPD and 120 SPD) but found a marked reduction (>50%) in protein and peptide coverage (**Figure 5A, B, Supplementary Table S5**). Interestingly, when comparing larger datasets of 40 SPD samples, 80 SPD, and 120 SPD, we found a large overlap between the methods, with only 451 proteins being unique to the 40 SPD runs (**Figure 5C**). These results suggest that comparisons between clusters of single cells might be similar when analyzed by 40 SPD and 120SPD. However, we show that many of the lowly-expressed proteins that are found robustly in 40 SPD runs, are only sporadically found in 80 and 120 SPD methods. As a result, robust clustering might not be feasible in these cases.

**Figure 5:**
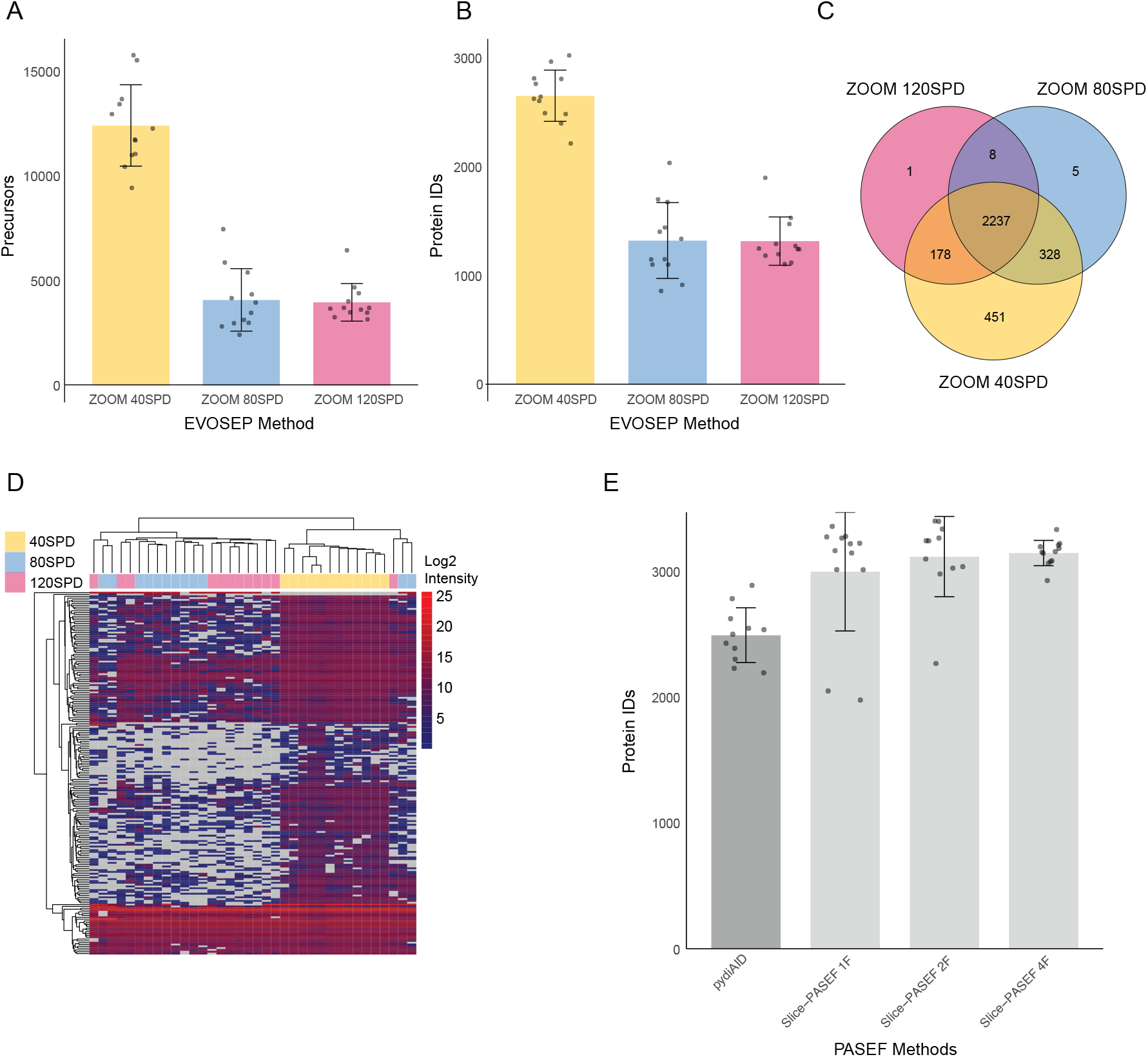
Optimization of LC-MS parameters for SCP. **A-B**. A Comparison of three Evosep1 LC methods determines the number of proteins (**A**) and precursors (**B**) identified using 40, 80, and 120 SPD methods. **C**. A Venn diagram shows the protein overlap of analyses of 36 cells analyzed by 40, 80, and 120 SPD methods. **D**. Hierarchical clustering of samples analyzed by each LC method shows similar expression levels of highly expressed proteins, but sporadic and inconsistent expression of lowly expressed proteins in 80 and 120 SPD, relative to 40 SPD. **E**. Examination of Slice-PASEF shows a significant advantage compared to the py_diAID optimized method. A minimum protein IDs cutoff was set to 1000 IDs.

In the next step, we wished to examine the impact of distinct TimsTOF SCP acquisition methods. In the last year, several approaches have been developed to ensure maximal coverage of the ion cloud. DIA-PASEF standard method was optimized by the py_diAID tool (25). We compared this standard method to the newly introduced Slice-PASEF method. Slice-PASEF employs continuous slicing of the precursor ion space with fragmentation of all ions in each slice (22). We optimized the method and tested 1-m/z frame, 2-frames, and 4-frames in each cycle. We found a marked improvement in protein identification using the Slice-PASEF method compared to DIA-PASEF, and we found only minor differences between the three Slice-PASEF options (**Figure 5E**).

### Single-cell analysis of tumor macrophages

Moving from cultured cells to in-vivo tumor cell populations poses additional challenges due to the smaller size of many cell types, particularly immune cell populations. We, therefore, investigated whether the optimized sample preparation protocol is sufficient to achieve biologically relevant information from immune cells in the tumor microenvironment. To that end, we injected MC38 murine cancer cells into the mouse tail vein to induce the development of lung metastases. After a week, we isolated the lungs and generated single-cell suspensions upon the appearance of small lesions. As a proof-of-concept, we focused our experiment on lung macrophages, sorted by flow cytometry and then dispensed as single cells into 1536-well plates on the CellenONE, as described above. Overall, we found slightly higher numbers of proteins/cell in the tumor macrophages compared to the control lung macrophages (injected with PBS). These reached a maximal number of proteins of more than 1800 proteins and 6000 precursors in a single cell (**Figure 6A, B, Supplementary Table S6**). Examination of the identity of these proteins showed that we can identify the most known macrophage markers, including CD68, MRC1 (CD206), ITGAX (CD11c), FCGR1 (CD64), etc. (**Figure 6C**). Furthermore, we were able to identify ∼500 significantly changing proteins between tumor macrophages and control lung macrophages (**Figure 6D**). Significant changes in such a large number of proteins already reflect that the current depth enables the identification of biologically relevant proteins. Specifically, we found that the tumor macrophages present increased expression of MHC molecules responsible for antigen presentation and multiple interferon-activated proteins (e.g., Ifi204, Ifi205, etc.). In agreement with the known anti-inflammatory role of MRC1 (26), we found higher levels of MRC1 in control macrophages than in tumor macrophages. These results show that with the current depth, we can capture the cellular activation state and associate these markers with hundreds of additional proteins in an unbiased way.

**Figure 6:**
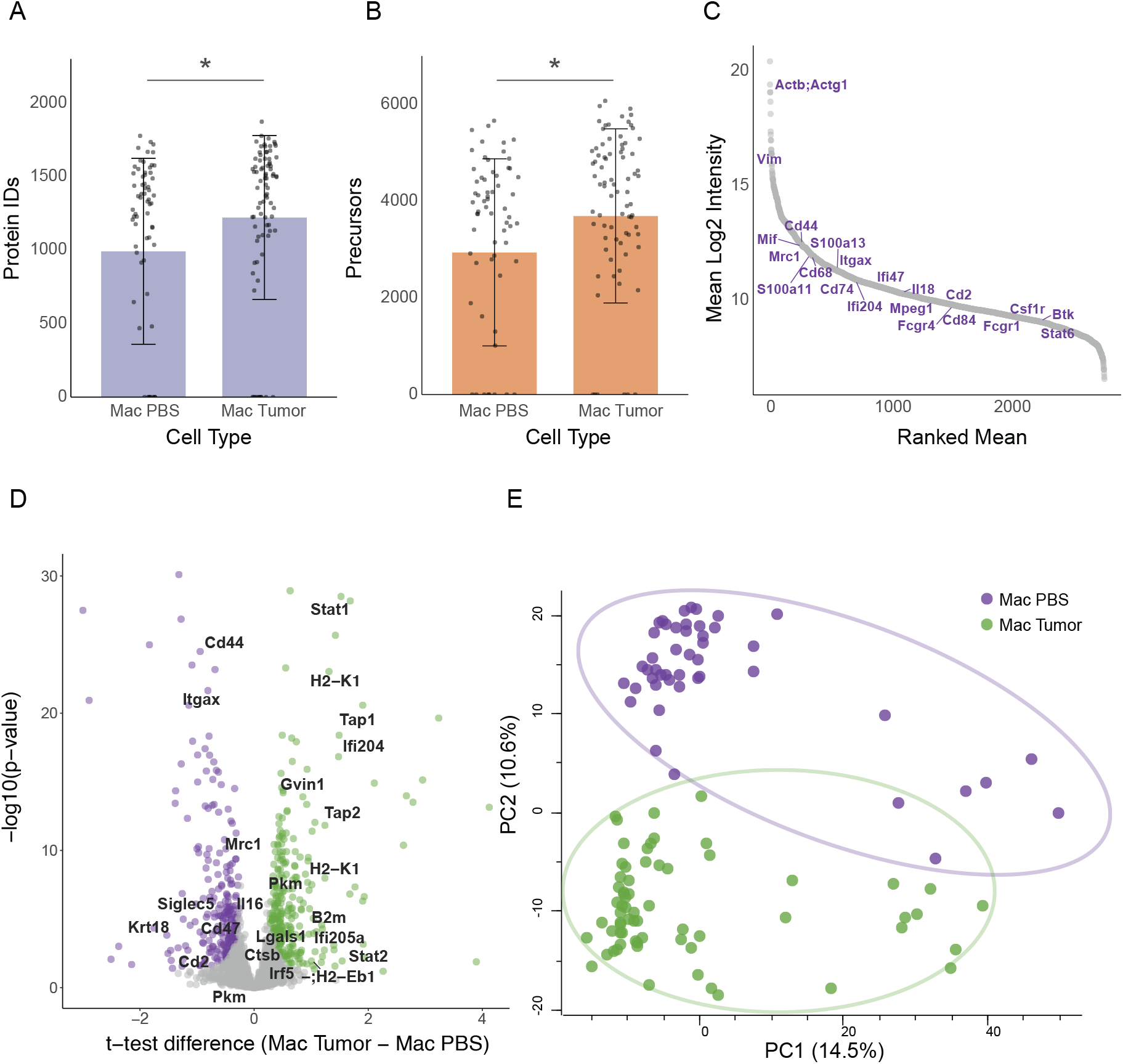
In-vivo macrophage SCP. **A-B**. Protein (**A**) and precursor (**B**) identification from lung macrophages from control lungs and tumor-bearing lungs. **C**. An aggregated S-plot of the average macrophage proteins identifies known macrophage markers. **D**. A volcano plot shows the significantly changing proteins between tumor and control macrophages (T-test with FDR= 5% and S_0_=0.1). **E**. Principal component analysis shows unsupervised separation between tumor and control macrophages.

Finally, we examined whether the current analysis enables unbiased separation between macrophage proteomes. PCA analysis separated the cells into two main clusters, which match the separation between tumor macrophages and controls (**Figure 6E**). Beyond the two main clusters, a few cells appeared as outliers. Analysis of the larger number of cells will suggest whether these can form additional smaller biologically relevant clusters. Overall, our results show a robust pipeline that enables the identification of important biological signals. We propose that this experimental approach can now be used for larger-scale SCP-based cancer research.

## Discussion

In this work, we describe a high-throughput sample preparation pipeline that enables a robust analysis of in-vivo cells. The 1536-well format provides a readily available platform that is combined with semi-automated Evotip loading and LC-MS analysis on the Evosep1-TimsTOF SCP using the SPD 40 method and Slice-PASEF. The coverage of ∼3000 proteins/cell is comparable to that of scRNAseq, but the low number of missing values is likely to simplify cell clustering and downstream statistical analyses. Notably, the experimental pipeline can also be combined with other instruments, and is not limited to the Evosep-TimsTOF setup.

The parallel processing of 1536 cells overcomes the first rate-limiting step of single-cell proteomics research. However, it transfers the limitation to the LC-MS acquisition step. We show that increasing the throughput to 120 SPD markedly reduces the number of identified proteins. Identification of ∼1500 proteins in HeLa cells using 120 SPD methods may be valuable. However, we speculate that the application of this method to in-vivo immune cells might not reach sufficient depth. To tackle the throughput challenges, several sample multiplexing methods have been proposed. MS-level multiplexing by plexDIA(27) or dimethyl labeling(28) provides accurate MS-level quantification. However, it increases throughput only 2-3fold. In contrast, TMT labeling provides higher throughput but complicates downstream analyses due to potential TMT-batch effects and ratio compression (29, 30). While some of these limitations are resolved in the analyses of single TMT batches, the limited depth of single-cell analyses is expected to result in complicated normalization challenges between TMT sets. Of note, high multiplexing with TMT can only be achieved with very high resolution that is not yet reached on the TimsTOF or the Astral analyzers(31).

Despite the limited LC-MS throughput, we show that the current state of single-cell proteomics already provides sufficient information to tackle biologically relevant questions in cancer biology and immunology. Tumor cells (cancer, stroma, and immune) can be used to unravel modes of cellular interactions and identify pathways related to cellular activation states. These studies are expected to provide important counterparts to the commonly used scRNAseq techniques. Ultimately, SCP will identify novel drug-resistance mechanisms and highlight potential cancer targets.

## Abbreviations

SCP: Single Cell Proteomics
MS: Mass Spectrometry
LC: Liquid Chromatography
PASEF: Parallel Accumulation Serial Fragmentation

## Acknowledgments

We thank the members of the Geiger lab for fruitful discussions and technical assistance. We thank Dr. Roni Oren and Mirie Zerbib from the Weizmann Institute Department of Veterinary Resources for their assistance in animal experiments and thank Alex Jahanfard from the Weizmann Institute Physics Core Facilities for designing and producing the 1536 adaptor. We thank the support of CellenOne, Agilent, Evosep and Bruker with the establishment of new protocols. This work was funded by the Israel Science Foundation grant #3495/19 and the European Council ERC-consolidator grant # 101044574.

